# Production, purification and evaluation of biodegrading potential of PHB depolymerase of *Stenotrophomonas* sp. RZS 7

**DOI:** 10.1101/698381

**Authors:** R. Z. Sayyed, S. J. Wani, Helal F. Al-Harthi, Asad Syed, Hesham Ali El-Enshasy

**Author notes:** **Corresponding author:** R.Z. Sayyed. These authors contributed equally to this work. These authors contributed equally to this work (Helal F. Al-Harthi, Asad Syed, Hesham Ali El-Enshasy).

## Abstract

There are numerous reports on PHB depolymerases produced by a wide variety of microorganisms isolated from various habitats, however, reports on PHB depolymerase isolated from plastic contaminated sites are scares. Thermophilic PHB polymerase produced by isolates obtained from plastic contaminated sites is expected to have better relevance for its application in plastic/bioplastic degradation. Although PHB has attracted commercial significance, the inefficient production and recovery methods, inefficient purification of PHB depolymerase and lack of ample knowledge on PHB degradation by PHB depolymerase have hampered its large scale commercialization. Therefore, to ensure the biodegradability of biopolymers, it becomes imperative to study the purification of the biodegrading enzyme system. We report the production, purification, and characterization of extracellular PHB depolymerase from *Stenotrophomonas* sp. RZS 7 isolated from a plastic contaminated site. The isolate produced extracellular poly-β-hydroxybutyrate (PHB) depolymerase in the mineral salt medium at 30oC during 4 days of incubation under shake flask condition. Purification of the enzyme was carried out by three different methods using PHB as a substrate. Purification of PHB depolymerase by ammonium salt precipitation, column chromatography, and solvent purification method was successfully carried out. Among the purification method tested, the enzyme was best purified by column chromatography on Octyl-Sepharose CL-4B column with maximum (0.7993 U mg^-1^ ml^-1^) purification yield. The molecular weight of purified PHB depolymerase (40 kDa) closely resembled with PHB depolymerase of *Aureobacterium saperdae*. Experiments on assessment of biodegradation of PHB in liquid culture medium and under natural soil conditions confirmed PHB biodegradation potential of *Stenotrophomonas* sp. RZS 7. The results obtained in FTIR analysis, HPLC study and GC-MS analysis confirmed the biodegradation attempt in liquid medium by *Stenotrophomonas* sp. RZS 7. Changes in surface morphology of PHB film in soil burial as observed in FE SEM analysis confirmed the biodegradation of PHB. The isolate was capable of degrading PHB and resulted in 87.74% degradation. Higher rate of degradation under natural soil condition is the result of activity of soil microbes that complemented the degradation by *Stenotrophomonas* sp. RZS 7.

## Introduction

Poly-β-hydroxy alkanoates (PHAs) or Poly-β-hydroxybutyrate (PHB) are accumulated as a source of food and energy by a wide variety of bacteria growing under nitrogen-deficient but conditions and is mobilized during nutrient stress under the influence of PHB depolymerase [1-3]. PHA/PHB is considered as the best eco-friendly and renewable alternative to the synthetics petrochemical plastics because of its similar properties to synthetic plastic [4-6] besides being thermoplastic and biodegradable in nature. Because of such useful properties, it has attracted commercial interest for use as the best alternative to the hazardous synthetic petrochemical polymers and hence it has been successfully commercialized [7-8]. During last decades much research has been devoted towards distribution and occurrence of PHB degraders and studies on different PHB depolymerases. Jendrossek and Handrick [9] reported that PHB depolymerases are responsible for extracellular PHB degradation. Extracellular PHB depolymerases of *Aspergillus fumigatus* Pdf1 [10], *A*. *Saperdae* [11], *Thermus thermophiles* HB8 [12], *Streptomyces bangladeshensis* 77T-4 [13], *Penicillium simplicissimum* LAR13 [14], *Acidovorax* sp. TP4 [15], *Emericellopsis minima* W2 [16] has been isolated and purified.

There are many different degradation mechanisms that are combined synergistically in nature to degrade polymers [17]. These mechanisms include bio-deterioration, bio-fragmentation, assimilation, mineralization and enzymatic degradation through extracellular and intracellular depolymerases that hydrolyze the polymer to water soluble products. PHA depolymerases being water soluble have an ability to bind specifically to polyester surfaces. Poly(HA)-degrading microorganisms secret specific poly(HA) depolymerases under starved conditions, which hydrolyze the polymer extracellularly to water soluble product and produce transparent clearing zones around the depolymerase secreting colonies (clear zone technique). Enzymatic hydrolysis of P(3HB) results in 3HB dimer as the major product besides small amounts of 3HB monomer. These cleave mainly the second and third ester linkages from the hydroxyl terminus [18].

However, the organism isolated from the plastic contaminated site and having the ability to degrade PHB may be a potential source of dynamic PHB depolymerase. However, inefficient production and recovery process of PHB, inefficient purification of PHB depolymerase and lack of ample knowledge on PHB degradation by PHB depolymerase have hampered the large scale commercialization of PHB. Therefore, to ensure the biodegradability of biopolymers, it becomes imperative to study the purification of biodegrading enzyme system [16].

The present paper reports the production, purification, and characterization of extracellular PHB depolymerase of *Stenotrophomonas* sp. RZS 7 isolated from a plastic contaminated site.

## Material and Methods

### PHB

All the experiment was carried out using PHB powder. PHB was obtained from Sigma-Aldrich, Germany.

### Source of culture

Fungal cultures such as *Aspergillus niger* NCIM1025, *Aspergillus flavus* NCIM650, *Alternaria alternate* ARI715, and *Cercospora arachichola were procured from* NCIM, NCL, Pune, Maharashtra, India. Bacterial cultres namely *Alcaligenes* sp RZS 4 and *Pseudomonas* sp. RZS 1 and actinomycetes culture namely *Streptomycetes s*p. were obtained from culture depository of the Department. *Stenotrophomonas* sp. RZS 7 isolated from plastic contaminated site was previously identified [19] and used in the present study as a source of PHB depolymerase.

### Production of PHB depolymerase

Production of PHB depolymerase was carried out at shake flask level by growing *Stenotrophomonas* sp. RZS 7 in minimal medium (MM) containing PHB, 0.15%; K_2_HPO_4_, 0.7 g; KH_2_PO_4_, 0.7 g; MgSO_4_, 0.7 g; NH_4_Cl, 1.0 g; NaNO_3_, 1.0 g; NaCl, 5 mg; FeSO_4_, 2 mg, ZnSO_4_, 7 mg in 1 L of distilled water [14] at 120 rpm for 4 days at 30°C.

### PHB depolymerase assay

Following 4 day’s incubation at 30°C and 120 rpm, MM broth was centrifuged a 10,000 rpm for 15 min and the supernatant was assayed for PHB depolymerase [12]. For this purpose PHB granule (substrate for PHB depolymerase) were sonicated (20 kHz for 15 min) and suspended in 50 mM Tris-HCl buffer (pH 7.0) and 150 μg ml^-1^ of this suspension and 2 mM CaCl_2_ was added in 50mM Tris-HCl buffer (pH 7.0) followed by the addition of culture supernatant (0.5 mL). Enzyme activity was spectrophotometrically measured at 650 nm as a decrease in the PHB turbidity. One unit of PHB depolymerase was defined as the quantity of enzyme required to decrease the absorbance by 0.1 min^-1^.

### Purification of PHB depolymerase

Having confirmed the presence of PHB depolymerase in cell-free supernatant of MM broth, the broth was subjected for purification of the enzyme by three approaches as follows

### Ammonium salt precipitation

The crude enzyme in the culture supernatant was precipitated by adding solid ammonium sulfate with continuous stirring at 4°C for 1 h. The precipitate was dissolved in the Tris-HCl buffer (pH 7) supernatant and dissolved precipitate was transferred in a separate dialysis bag and allowed for overnight dialysis in chilled phosphate buffer [13]. The protein concentration of dialyzed supernatant, as well as dialyzed precipitate, was measured by the Lowry method [20]. PHB depolymerase activity of dialyzed supernatant and dialyzed precipitate was measured as described earlier.

### Solvent purification method

The culture supernatant of the isolate was centrifuged at 10,000 rpm for 20 min. The residue obtained was dissolved in pre-chilled 1:1 acetone ethanol mixture, shaken well and kept in a water bath at 50°C until all solvent is evaporated. The pellet obtained after evaporation was dissolved in Tris-HCl buffer (pH 7). PHB depolymerase activity of pellet, as well as supernatant, was carried out as described.

### Column chromatography

The cell-free supernatant of the isolate was applied onto an Octyl-Sepharose CL-4B column pre-equilibrated with 50 mM glycine NaOH buffer (pH 9.0) and eluted with a gradient of 0 to 50% ethanol [12], the fractions were collected and PHB depolymerase activity of each fraction was determined as described.

### Determination of molecular weight of purified PHB depolymerase

To measure the molecular weight of purified PHB depolymerase, SDS PAGE (Sodium Dodecyl Sulfate Polyacrylamide Gel Electrophoresis) was performed with standard molecular weight markers such as phosphorylase b (82.2 kDa), bovine serum albumin (64.2 kDa), egg albumin (48.8 kDa), carbonic anhydrase (37.1 kDa), trypsin inhibitor (25.9 kDa), lysozyme (19.4 kDa), lysozyme (14.8 kDa) and lysozyme (6.0 kDa). The protein concentration of purified band was measured by the Lowry method with bovine serum albumin (BSA) as a standard [14].

### Assessment of biodegradation of PHB

To evaluate the efficacy of degradation of biopolymer, following methods were used

### PHB biodegradation under laboratory conditions

#### Preparation of PHB film

Preparation of PHB film was done as per the method of Liang et al. [21]. For this purpose PHB powder (0.15 gm) was dissolved in 50 ml chloroform and vortexed for 10-15 min. The solution was poured into clean, dry Petri dishes and kept overnight at room temperature. PHB film was obtained following the evaporation of chloroform. This film was used for assessment of biodegradation.

PHB film was separately added as a sole carbon source into MSM and inoculated with *Stenotrophomonas* sp. RZS 7 and incubated at 30oC for 10 days at 120 rpm and observed for degradation of PHB film. Qualitatively estimation involved the weight loss method. Quantitatively estimation involved the measurement of change in optical density and crotonic acid concentration (μg mL^-1^) at different time interval [22] which is based on the conversion of PHB to crotonic acid when heated in presence of concentrated sulphuric acid. The absorbance was measured at 235 nm and the amount of crotonic acid was calculated from calibration curve prepared with standard crotonic acid in the range of 10-100 μg mL^-1^.

### PHB biodegradation under natural soil conditions

To check the biodegradability and durability of polymer sample under natural soil conditions, soil burial test method was carried out. Soil burial test methods are used to give an indication of the duration of polymer in a given soil under specific conditions. To study the biodegradation of PHB film, garden soil was taken in different trays (1800 gm per tray) and the polymeric film covered with net was incorporated into the tray with respective inoculum. The trays were covered with moist thick paper [23]. Degradation of PHB film by *Stenotrophomonas* sp. RZS 7 was observed in five different sets and was compared with negative control containing only sterile soil. The degradation of PHB film was expressed as percent reduction in the weight of PHB film using following formula

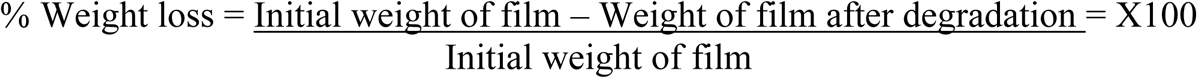

### Fourier-Transform Infrared Spectroscopy (FTIR) analysis

Following the desired incubation period for degradation, the inoculated and incubated liquid culture medium of *Stenotrophomonas* sp. RZS 7 was subjected for FTIR analysis by using FTIR (FTIR Model 8400S, Shimadzu, Japan,) using IR Solution version 1.40 software. FTIR spectra were recorded in the range of 4000 to 500 cm^-1^ [24]. The functional groups present in degradation pattern were determined by FTIR and compared with the control.

### High Performance Liquid Chromatography (HPLC) analysis

For HPLC analysis of PHB film, the content compound was analyzed by HPLC (Younglin, South Korien, ACME-9000) [25]. For this purpose 20 μl of sample was injected into the C18 column (4.6 × 250 mm, particle size 5 µm) as stationary phase and allowed to run on mobile phase consisting of acetonitrile : water (30:70 v/v) at 254 nm with flow rate of 0.6 mL min^-1^. The peaks were analyzed by using Autochro-3000 software.

### Gas Chromatography-Mass Spectroscopic (GC-MS) analysis

#### Sample preparation

The liquid culture supernatant (2 mL) which was subjected for biodegradation by *Stenotrophomonas* sp. RZS 7 mixed with 2 ml chloroform and 2 ml of acidified methanol (acidified with 3% (v/v) H_2_SO_4_). For depolymerization and methanolysis of PHB the mixture was heated at 100°C for 3 h. During heating the mixture was shaken occasionally and subsequently cooled to room temperature. After heating at 100°C for 3 h, 1 ml of distilled water was added to the reaction mixture and the mixture was then vortexed for 2 min and allowed to stand for 10 min to separate the phases. At the bottom layer organic phase was obtained by glass Pasteur pipette and it was subjected for further analysis [26].

#### GC-MS analysis

For identification of monomers present in degraded sample, a coupled gas chromatography/mass spectrophotometry (GC-MS) was performed using gas chromatograph (Perkin Elmer, USA, Model Autosystem XL) with turbomass GC+ equipped with an SGE forte GC capillary column BP20 (30 m X 0.250 μ internal diameter X 0.25 μ) (Australia) and mass spectrophotometer. The mass spectra obtained were compared with the spectra of standard methyl esters of PHB available from National Institute of Standards and Technology (NIST) mass spectra library. The peak identities were first confirmed by matching relative retention time with respect to standard sample extracted at time zero.

#### GC-MS parameters

One μL of PHB degrading sample was injected by split injection with a split ratio of 10:1 using 10 μL syringe (SGE analytical, Australia). Helium was used as carrier gas at a flow rate of 1 mL min^-1^. The oven temperature of column was programmed from 120°C for 2 min, increased at a rate of 20°C min^-1^ to 230°C, and held at this temperature for 10 min. The temperature of injector and detector was 225°C and 230°C respectively.

### Field Emission Scanning Electron Microscopy (FE SEM) analysis

The surface morphology of the PHB film was analyzed through Field Emission Scanning Electron Microscopy (FE SEM, Model S-4800, Hitachi, Japan) to check any structural changes following PHB degradation in soil burial experiment. After throughout washing with sterile distilled water, samples were mounted on stub using carbon tape. Gold coating was carried out in vacuum by evaporation in order to make the sample conducting. Polymer film was dried in Infrared (IR) lamp. The images of the test samples were compared with those recorded on the original untreated samples.

### Statistical analysis

All the experiments were performed in triplicate and the mean of three replicates was considered. Each mean value was subjected to Student’s *t-*test and values of *P* ≤ 0.05 were taken as statistically significant [27].

## Results and Discussion

### Production of PHB depolymerase

After 4 days’ incubation at 30°C at 120 rpm in MM, *Stenotrophomonas* sp. RZS 7 yielded 0.721 U mL^-1^ PHB depolymerase.

### Purification of PHB depolymerase

#### Ammonium salt precipitation

With increasing concentrations of ammonium sulfate in the cell-free culture supernatant of *Stenotrophomonas* sp. RZS 7, increased precipitation of protein was observed, maximum proteins were precipitated with 70% ammonium sulfate concentration (Table 1). The protein concentration and PHB depolymerase specific activity and enzyme activity of dialyzed precipitate of *Stenotrophomonas* sp. RZS 7 were 0.219 mg mL^-1^, 0.7031 U mg^-1^ mL^-1^ and 0.154 U mL^-1^ respectively. Zhou et al. [28] have reported precipitation of PHB depolymerase of *Escherichia coli*, and *Penicillium* sp. DS9701-D2 by using 70 and 75% of ammonium sulfate Shivkumar et al. [21] have reported efficient precipitation of PHB depolymerase of *Penicillium citrinum* S2 with 80% of ammonium sulfate.

**Table 1.** Purification of PHB depolymerase of *Stenotrophomonas* sp. RZS7 on Octyl sepharose column

| Fraction | Total activity (U mL<sup>-1</sup>) | Total protein (mg mL<sup>-1</sup>) | Specific activity (U mg<sup>-1</sup>/mL<sup>-1</sup>) | % activity |
| --- | --- | --- | --- | --- |
| 1 | 0.027 (0.027) | 0.031 (0.010) | 0.0782 (0.032) | 11.13 (0.012) |
| 2 | 0.101 (0.011) | 0.153 (0.037) | 0.3781 (0.041) | 48.31 (0.039) |
| 3 | 0.247 (0.015) | 0.309 (0.074) | 0.7993 (0.090) | 79.93 (0.092) |
| 4 | 0.017 (0.001) | 0.015 (0.009) | 0.0106 (0.011) | 04.01 (0.012) |
| 5 | 0.003 (0.001) | 0.006 (0.005) | 0.0011 (0.008) | 00.10 (0.002) |

Values were taken to be statistically significant at $P \leq 0.05$
Values are the mean of three replicates

### Solvent purification method

Solvent purification of PHB depolymerase of *Stenotrophomonas* sp. RZS 7, retained only 51.14 % and 38.81% enzyme activity in pellet and supernatant (Table 2).

**Table 2.** – Purification profile of PHB depolymerase by various methods.

| Purification method | Enzyme activity (U mg<sup>-1</sup> ml<sup>-1</sup>) |
| --- | --- |
| Salt precipitation | 0.7031 (0.090) |
| Solvent purification | 0.1120 (0.022) |
| Column chromatography | 0.7993 (0.099) |

Values were taken to be statistically significant at $P \leq 0.05$
Values are the mean of three replicates

The protein concentration and PHB depolymerase specific activity and enzyme activity in pellet and supernatant were 0.219 mg mL^-1^, 0.5114 U mg^-1^ mL^-1^ and 0.3881 UmL^-1^ respectively. This significant loss of enzyme activity may be due to precipitation resulting in denaturation of proteins by solvent system. Thus the solvent purification method proved insignificant vis-à-vis other purification methods.

### Column chromatography

The dialyzed precipitate of *Stenotrophomonas* sp. RZS 7 obtained after ammonium salt precipitation when applied onto an Octyl-Sepharose CL-4B column pre-equilibrated with 50 mM glycine NaOH buffer (pH 9.0) and eluted with ethanol, yielded 5 fractions. Among all the fractions analyzed for PHB depolymerase enzyme activity, 3^rd^ fractions showed maximum enzyme activity (Table 1). PHBV depolymerase of *Bacillus* sp. AF3 and *Streptoverticillium kashmirense* AF1 have also been purified on Sephadex G-75 (30). Kim et al. [31] have also purified PHB depolymerase of *Emericellopsis minima* W2 and *Streptomyces* sp. KJ-72 on Sephadex G-100 and Sephadex G-150 respectively. Papaneophytou et al. [12] and Hsu et al. [13] have isolated and purified extracellular PHB depolymerases of *Thermus thermophiles* HB8 and *Streptomyces bangladeshensis* 77T-4 respectively using column chromatography.

Among all three purification methods, purification on Octyl sepharose CL-4B column gave good purification yield (Table 1), more enzyme activity and more specific activity. The protein concentration and PHB depolymerase specific activity and enzyme activity were 0.247 mg mL^-1^, 0.7993 U mg^-1^ mL^-1^ and 0.309 U ml^-1^ respectively.

### Determination of molecular weight of purified PHB depolymerase

The protein fraction of *Stenotrophomonas* sp. RZS 7 obtained from column chromatography having maximum PHB depolymerase enzyme activity showed single protein band in SDS PAGE having a molecular mass of approximately 40 kDa lying between molecular weight markers of 37.1 and 48.8. Sadocco et al. [11] have also reported the molecular mass of PHB depolymerase from *Aureobacterium saperdae* ranges from 42.7 kDa analyzed by SDS PAGE.

### Assessment of biodegradation of PHB

Among the various reference cultures taken from our own departmental culture depository all fungal and bacterial cultures were unable to degrade PHB when grow in MSM containing PHB as the only carbon source. Only *Streptomycetes* sp. was found to degrade PHB on 10th day of incubation but the extent of degradation was less. Hence for further biodegradation studies *Stenotrophomonas* sp. RZS 7 was used.

### Degradation of PHB film

#### Qualitative measurement

A gradual reduction in the weight of PHB film was recorded after 8 days of incubation fragmentation in PHB film was observed whereas no changes observed in control sample of PHB film (Table 3). Varda et al. [32] have also reported similar findings in case of microbial degradation of starch based plastic by *S*. *epidermidis*.

**Table 3.**
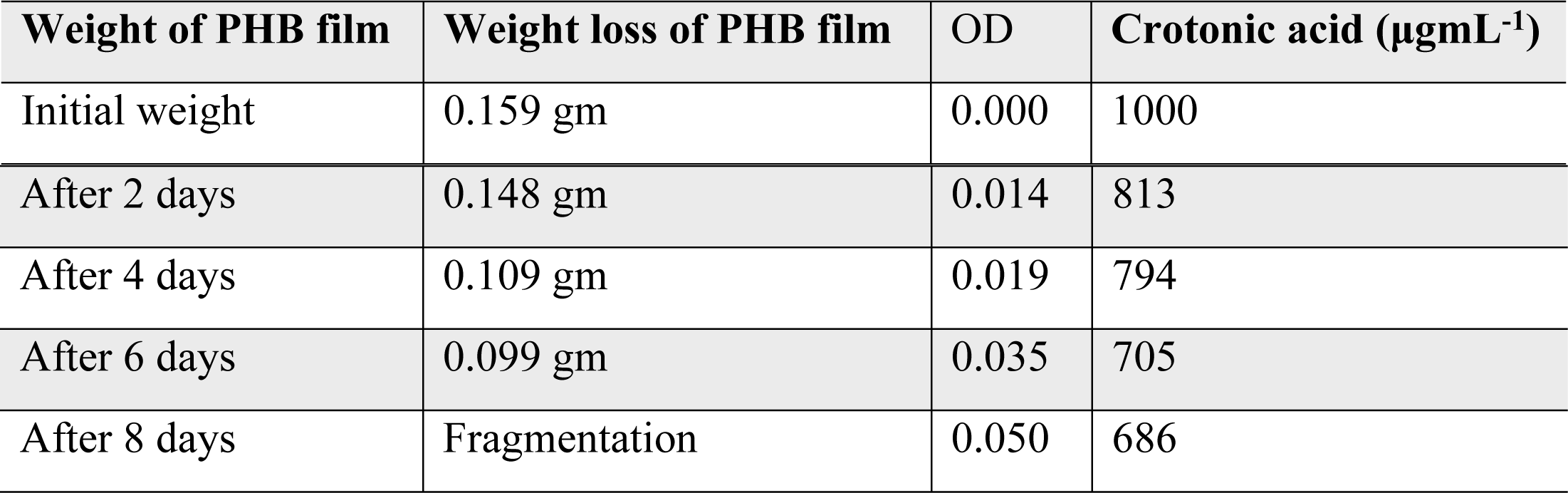
Qualitative measurement of biodegradation of PHB film

#### Quantitative estimation

Isolate RZS7 caused the reduction in the amount of crotonic acid with increase in the biomass. Maximum reduction in the amount of crotonic acid was recorded on 10th day of incubation at 37°C (Table 3). The reduction in crotonic acid concentration can be taken as the evidence of biodegradation of PHB film.

### Fourier Transform Infrared Spectroscopy (FTIR) analysis

FTIR analysis of liquid culture medium containing PHB film and inoculated with *Stenotrophomonas* sp. RZS 7 showed changes in the functional groups and significant shift of wave numbers, indicating the biodegradation of polymer. It was observed that in case of control preparation (without inoculum) FTIR chromatogram peaks appearing at 3020.63 cm^-1^ changed to 3018.70, 2926.11, 2856.67 cm^-1^, 2399.53 cm^-1^ changed to 2360.95, 2333.94 cm^-1^, 2088.98 cm^-1^ changed to 2079.33 cm^-1^, 1641.48 cm^-1^ shifted to 1629.90 cm^-1^ and 1631.83 cm^-1^ shifted towards 1529.60, 1427.37, 1381.08 cm^-1^ (Fig.1 and 2). The major functional groups in control sample were found at wavelength frequency 1641.48 and 1631.83 denoted C=O stretch in ketone, frequency 3020.63 denoted C-H which stretch in alkane, wavelength frequency 1014.59 denoted C-O which stretch in ester and 756.12 denoted C-H which is rock alkane. However the major functional group frequency changes observed in degrading sample i.e. 1629.90 denoted C=O which stretch in ketone, 1215.19 which denote alkyl halide, 2399.53,2360.95 and 2333.94 denoted H-C=O group, 927.79 denoted C-O which stretch in ester, 769.62 denoted C-H which is rock alkane. These changes in wavelength frequency of major functional group in degrading sample as compared to control give preliminary proof of biodegradation of polymer by *Stenotrophomonas* sp. RZS 7.

**Figure 1.**
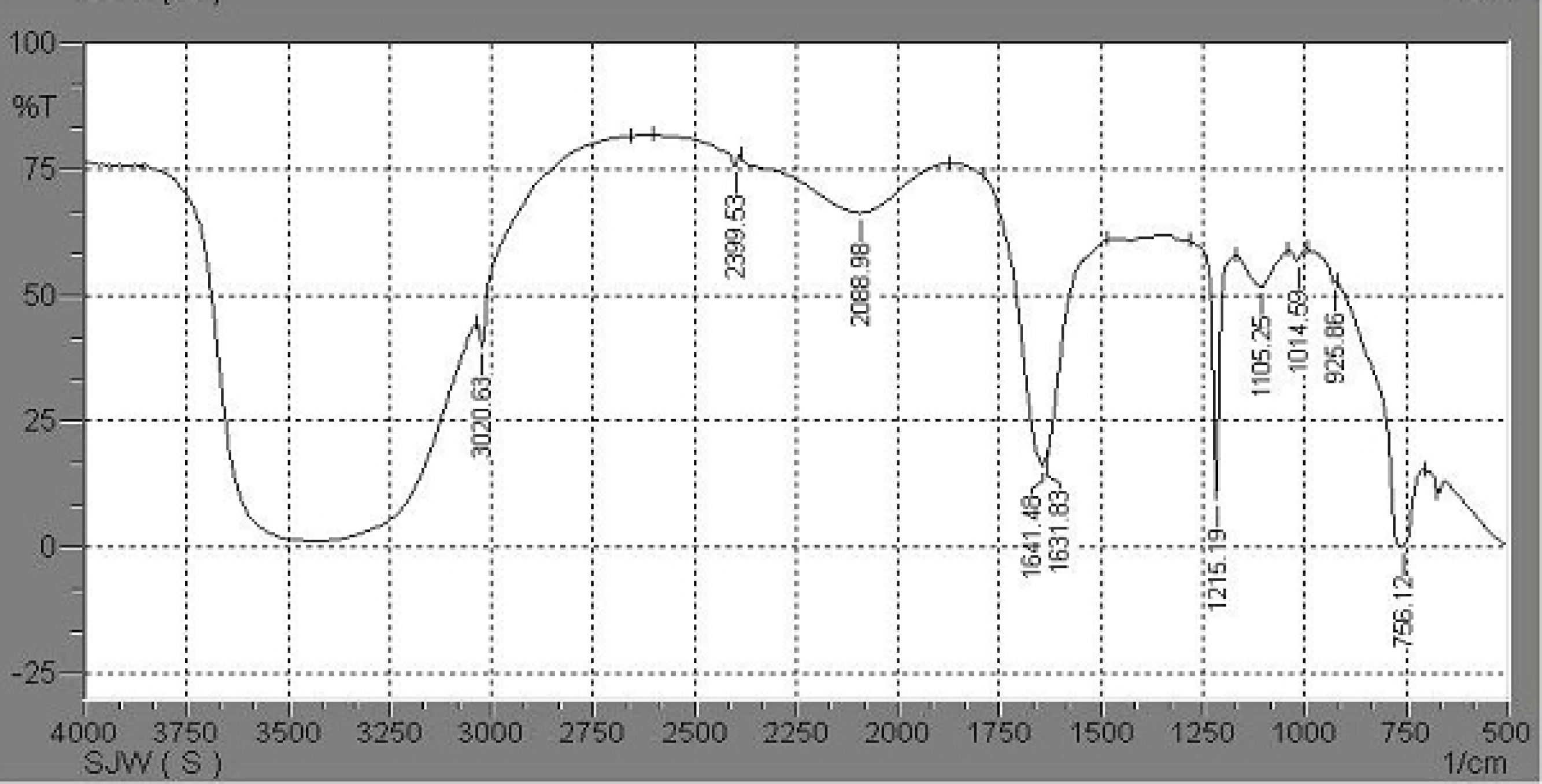
FTIR chromatogram of PHB (Control) carried out by using IR solution version 1.40 software. Spectra recorded in the range of 4000 to 500 cm^-1^.

**Figure 2.**
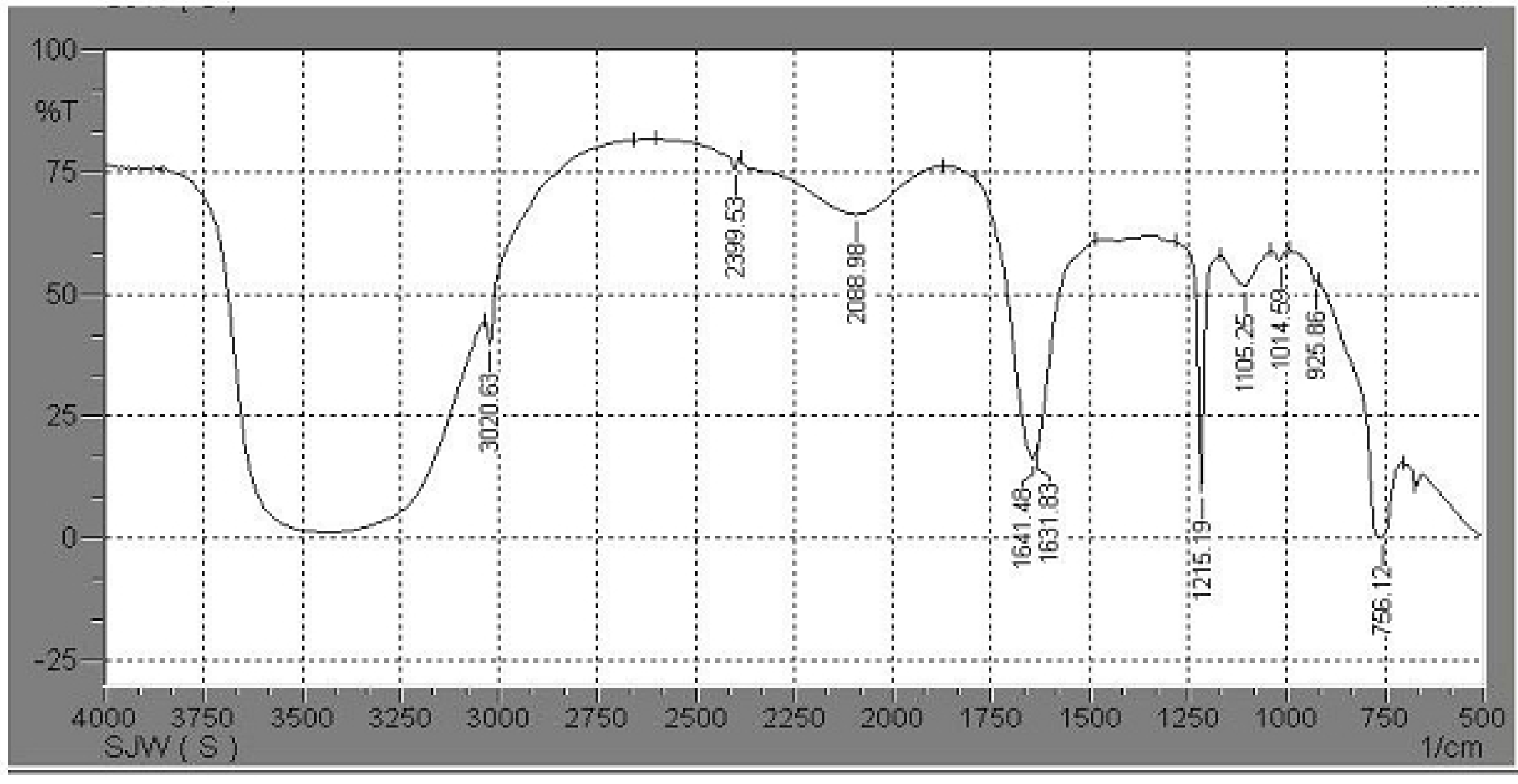
FTIR chromatogram of PHB after degradation by *Stenotrophomonas* sp. RZS 7 carried out by using IR solution version 1.40 software. Spectra recorded in the range of 4000 to 500 cm^-1^. The functional groups present in the spectra were compared with the functional groups in control preparation.

### High Performance Liquid Chromatography (HPLC) analysis

The spectra of the original polymer sample before degradation revealed six peaks at 254 nm during 20 min flow at the rate of 0.6 mL min^-1^ (Fig. 3). While polymer sample degradation by *Stenotrophomonas* sp. RZS 7 revealed nine peaks having different retention time as compared to control (Fig. 4) indicating the formation of monomers. The sum of total area of peaks of degraded sample was less as compared to the sum of total area of peaks of control (undegraded) preparation. This reduction in the peak area confirmed the degradation of polymer.

**Figure 3.**
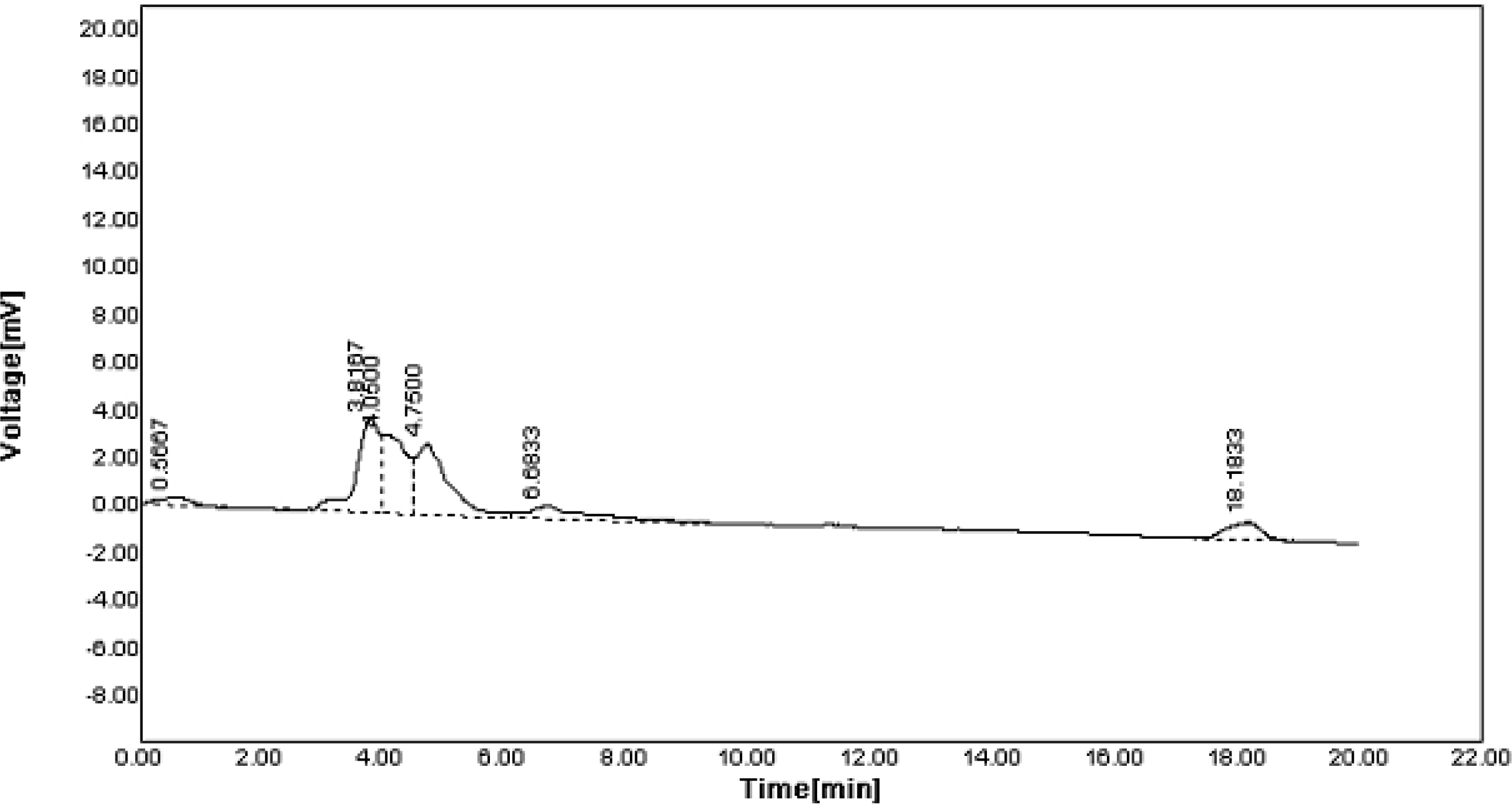
HPLC spectra of polymer before degradation (Control) recorded at 254 nm run on C18 column with mobile phase of acetonitrile : water (30:70 v/v) with flow rate of 0.6 mL min^-1^. Analysis was carried out at 254 nm with flow rate of 0.6 mL min^-1^. The peaks were analyzed by using Autochro-3000 software.

**Figure 4.**
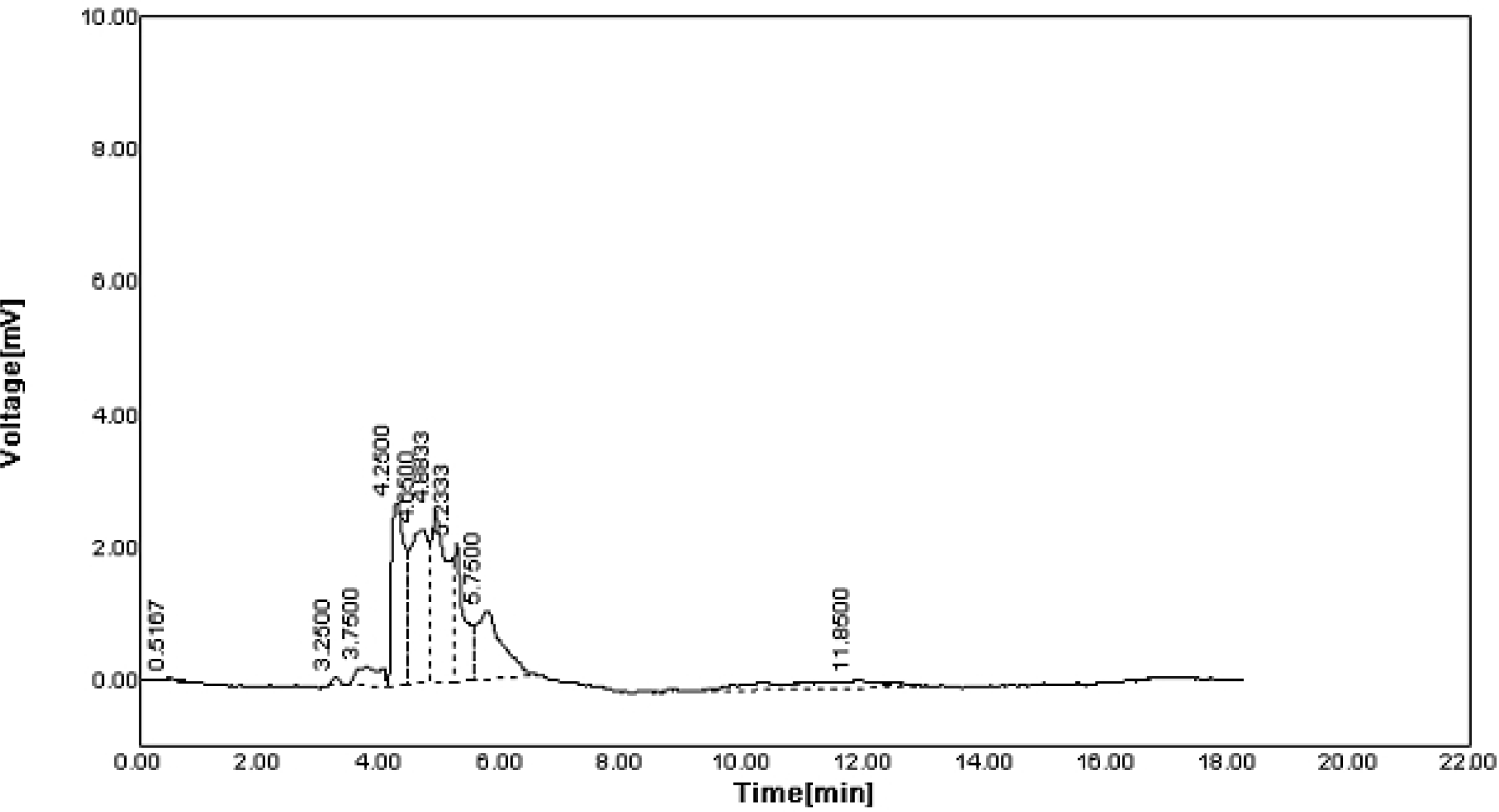
HPLC spectra of polymer after degradation recorded at 254 nm run on C18 column with mobile phase of acetonitrile : water (30:70 v/v) with flow rate of 0.6 mL min^-1^. The peaks were analyzed by using Autochro-3000 software and compared with the peaks of control.

### Gas Chromatography-Mass Spectroscopic (GC-MS) analysis

The degradation pattern on the basis of removal or decrease in the retention time of the peaks and their comparison with retention time of the peaks of standard demonstrated that the disappearance of peaks indicating degradation of PHB components by *Stenotrophomonas* sp. RZS 7 (Fig. 5 and Fig. 6). Significant decrease in area height and percentage in degraded sample was clear over the control (undegraded) sample. The total area of peaks of degraded sample was less as compared to the total area of peaks of control (undegraded) preparation. This reduction in the peak area confirmed the degradation of polymer into monomers.

**Figure 5.**
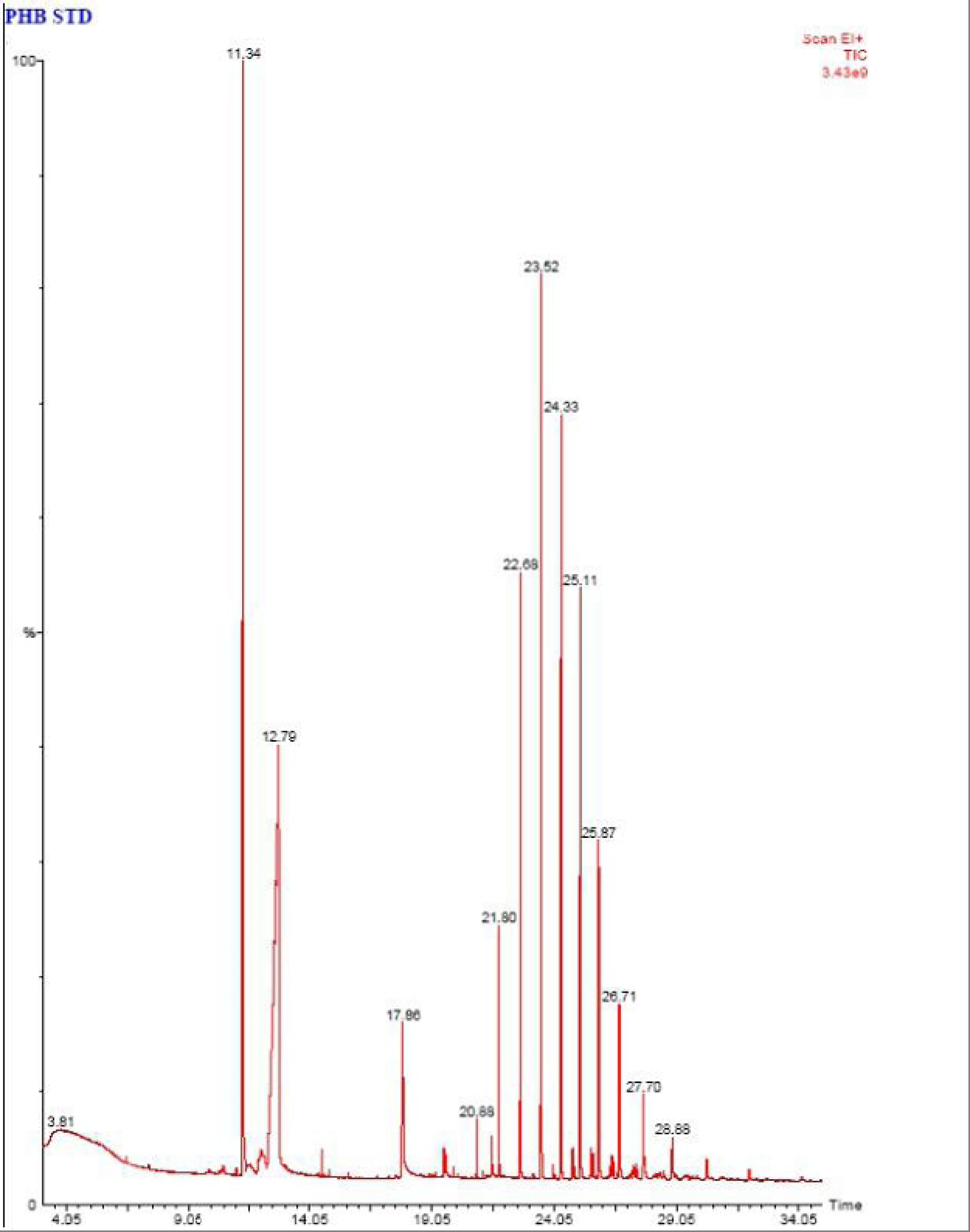
-Chromatogram of polymer sample before biodegradation (Standard). GC MS analysis was performed on gas chromatograph equipped with capillary column BP20 (30 m X 0.250 μ internal diameter X 0.25 μ) and mass spectrophotometer. The mass spectra obtained were recorded.

**Figure 6.**
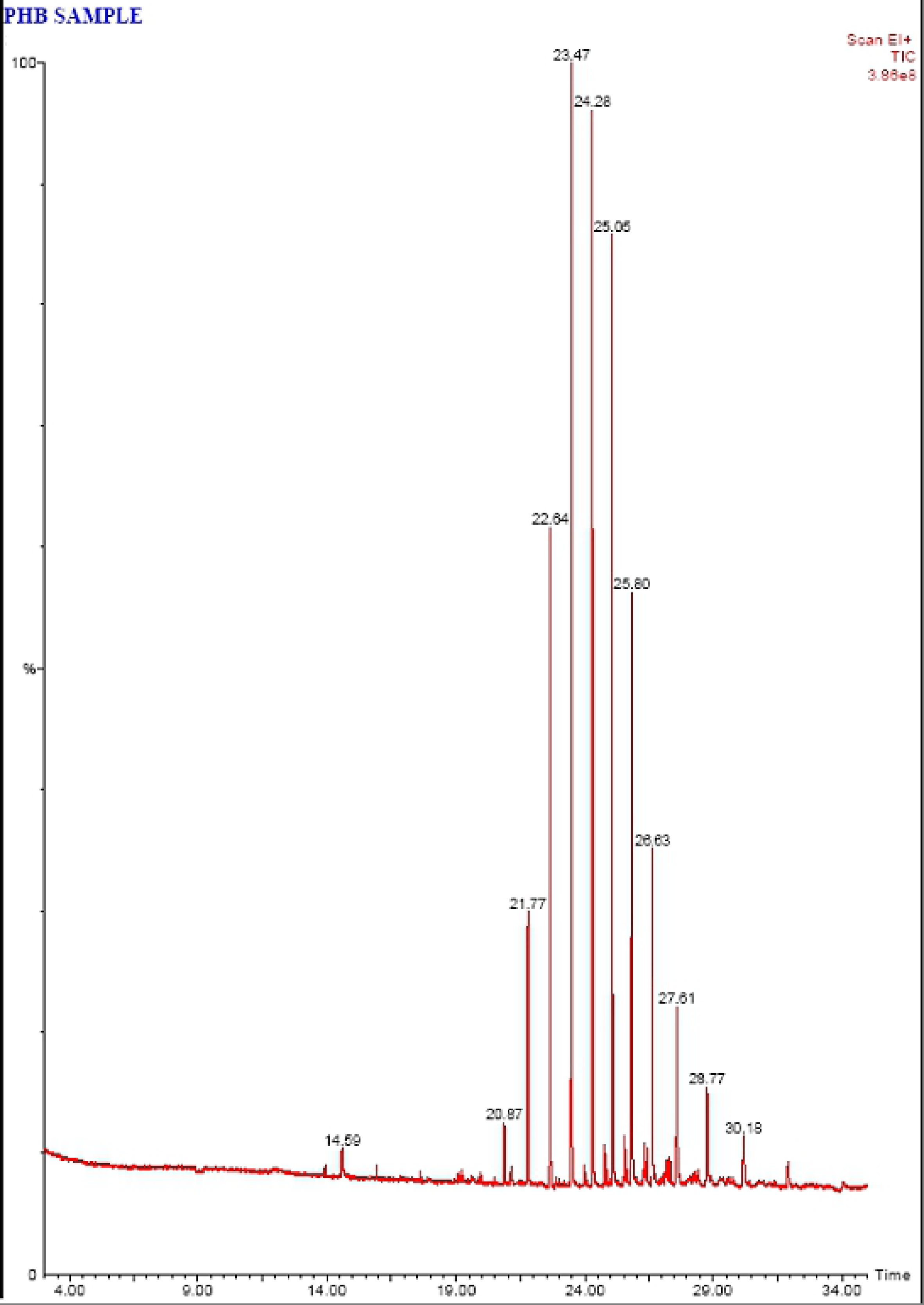
-Chromatogram of polymer sample after biodegradation. GC MS analysis was performed by using gas chromatograph with turbomass GC capillary column BP20 (30 m X 0.250 μ internal diameter X 0.25 μ) and mass spectrophotometer. The mass spectra obtained were compared and identified with the library of spectra of standard methyl esters of PHB. The peak were identified by matching relative retention time with respect to standard sample extracted at time zero.

The analysis of potential fragmentation patterns with GC-MS and the identities of specific peaks in the mass spectra were correlated to the carbonyl and hydroxyl ends of the represented hydroxyalkanoates ethyl esters of PHB after degradation. Tridecanoic acid 12-methyl-methyl ester and heptacosanoic acid methyl ester in standard polymer sample were completely degraded whereas Decanoic acid methyl ester, undecanoic acid methyl ester and nonanoic acid methyl ester were detected in degraded sample (Table 4). McLafferty and Turecek (33) reported that the mass spectra of the methyl esters of PHB dominated by m/z = 74 represented the carbonyl end of the molecule and showed the characteristic of 3-hydroxyl functional groups. However the peak at m/z = 59 represented the hydroxyl end released when molecule was cleaved at the bond between C3 and C4.

**Table 4.**
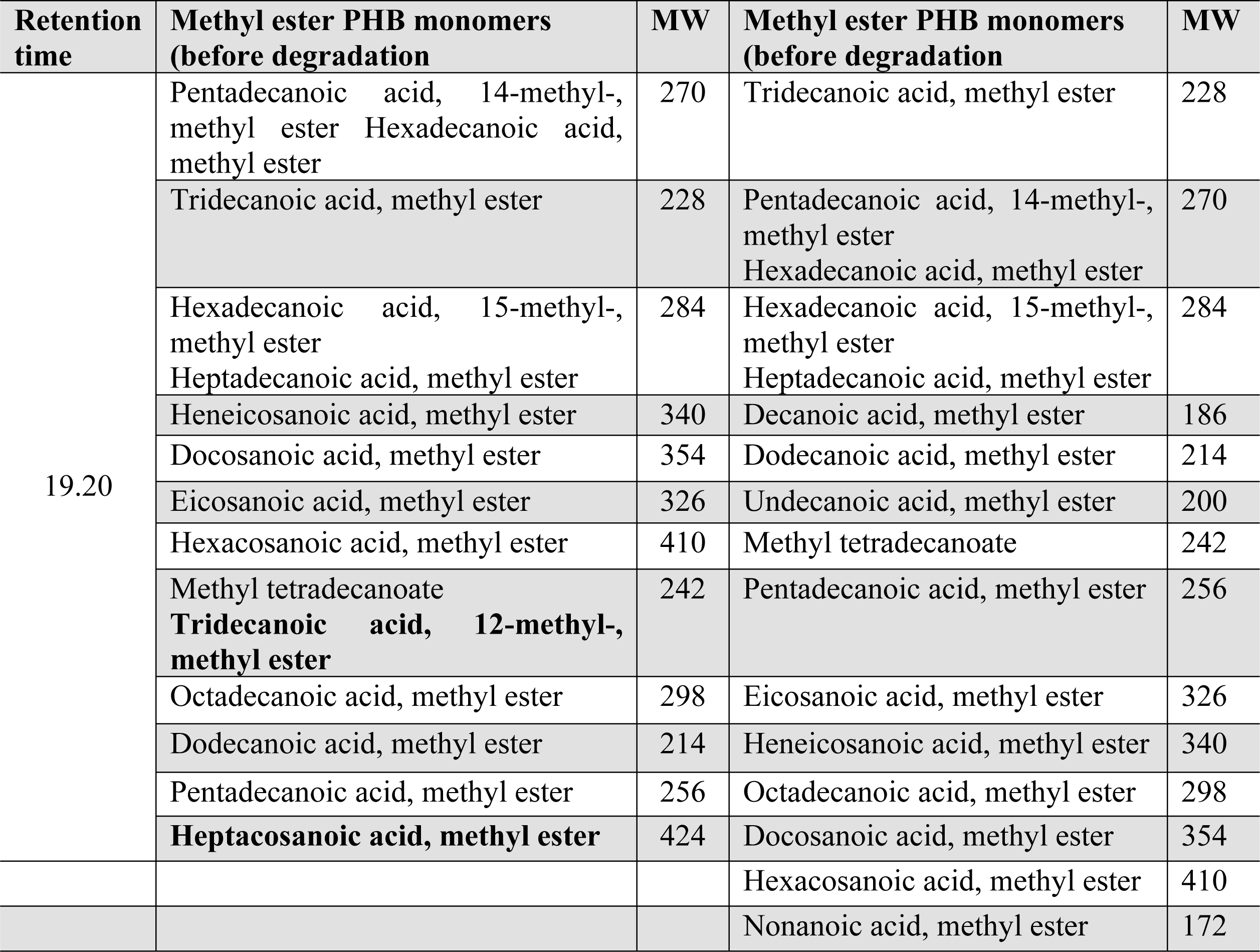
– GC MS analysis of methanolysed polymer before degradation (Standard) and after degradation

The mass spectra of standard polymer sample and the characteristics signals were recorded at m/z = 29, 39, 41, 43, 45, 55, 57, 59, 74, 75, 87, 101, 111, 131, 143, 157, 171, 185, 199, 219, 227, 239, 265, 270, 271, 314, 415. However in polymer degrading sample m/z = 29, 32, 41, 43, 44, 55, 57, 69, 74, 75, 87, 97, 111, 129, 143, 145, 169, 176, 199, 220, 227, 270 (Fig. 7) which can be attributed to oligomers of PHB (Fig. 8). Wennan et al. [34] reported that m/z = 29, 32, 41, 43, 55, 69, 75, 87, 97, 111, 143 denotes Pentadecanoic acid methyl ester, m/z = 29, 41, 55, 57, 69, 74, 75, 87, 97 denotes Decanoic acid, methyl ester, m/z = 41, 43, 55, 74 denotes Dodecanoic acid methyl ester, m/z = 74 denotes Hexadecanoic acid methyl ester, m/z = 43, 55, 74 denotes Octadecanoic acid methyl ester.

**Figure 7.**
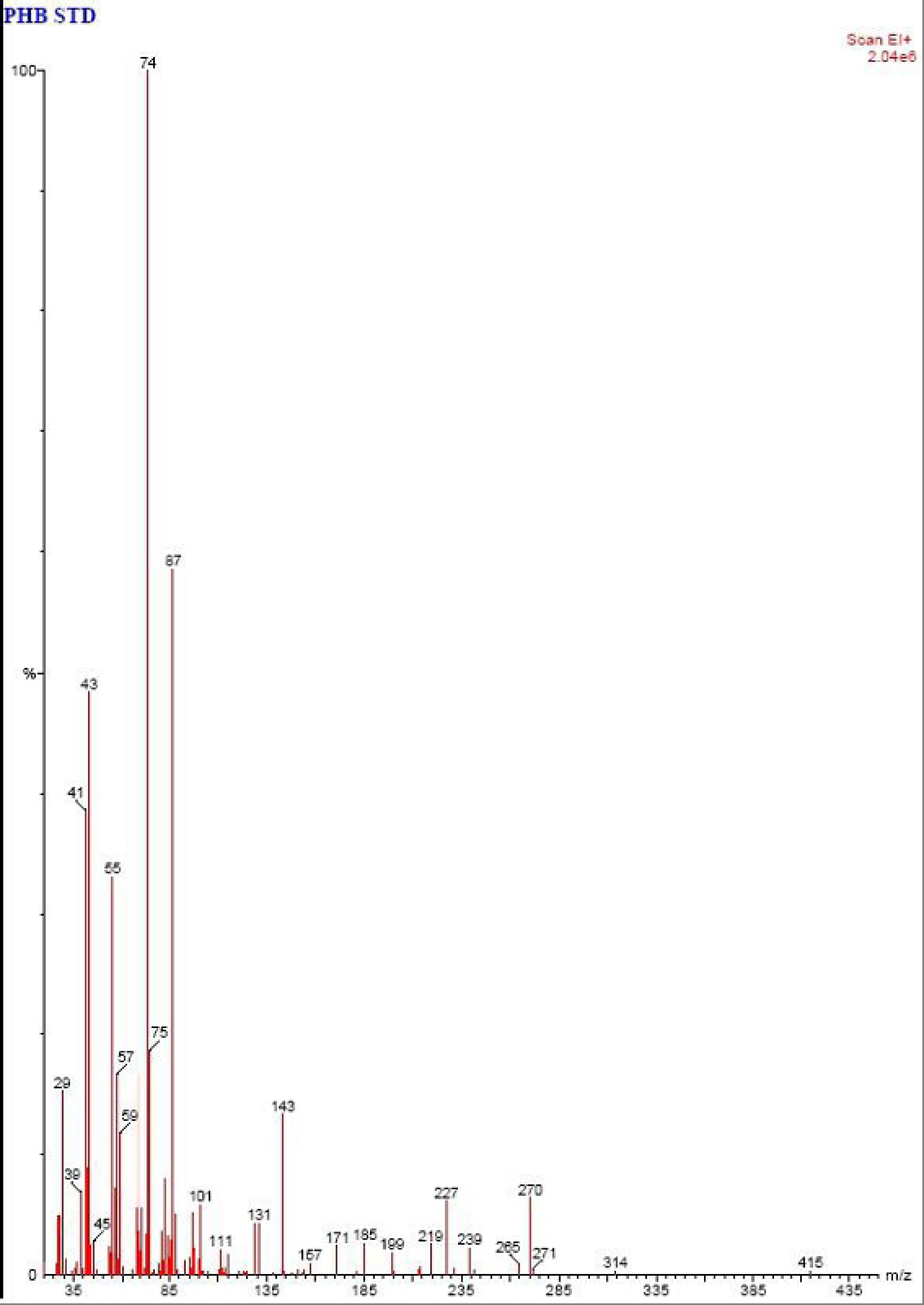
-GC-MS spectra of polymer sample before degradation showing mass spectra of m/z = 29, 39, 41, 43, 45, 55, 57, 59, 74, 75, 87, 101, 111, 131, 143, 157, 171, 185, 199, 219, 227, 239, 265, 270, 271, 314, 415 indicating the presence of intact polymer of PHB.

**Figure 8.**
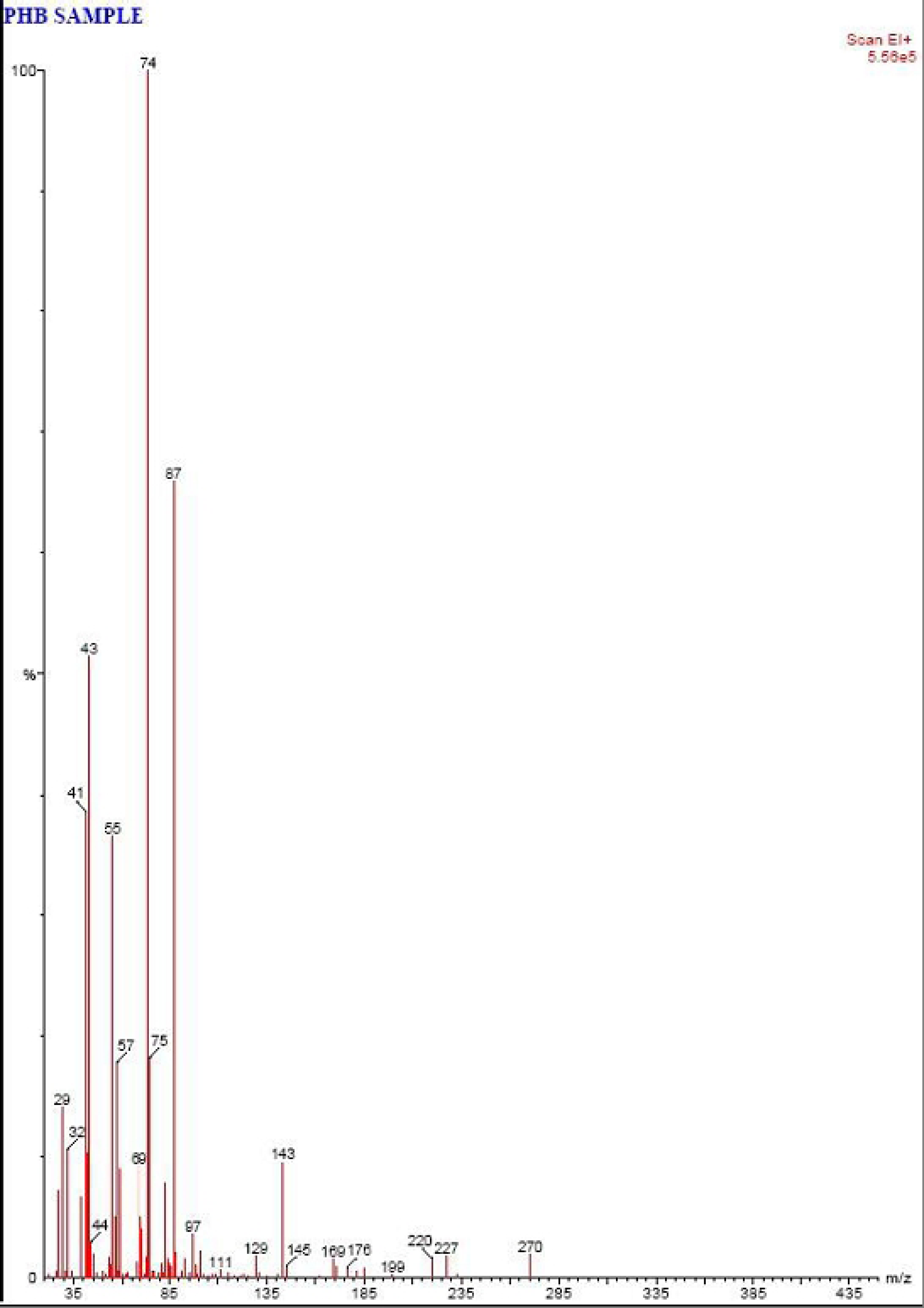
-GC-MS spectra of polymer sample after degradation showing mass spectra of m/z 29, 32, 41, 43, 44, 55, 57, 69, 74, 75, 87, 97, 111, 129, 143, 145, 169, 176, 199, 220, 227, 270 indicating the presence of oligomers of PHB.

### Field Emission Scanning Electron Microscopy (FE SEM) analysis

Surface analysis of PHB film after soil burial experiment showed many differences in the morphology of PHB film as compared to control (Figure 9A) and PHB film buried in soil (Figure 9B). Biodegradation of PHB film caused erosion and roughening of surface of PHB film (Figure 9C). Gautam and Kaur [35] have also reported the morphological changes in polyethylene surface after degradation compared with control following SEM analysis. Calabia and Tokiwa [36] analyzed the degradation of PHB by scanning electron microscopy, and observed the presence of hemispherical holes on the film surface after degradation and claimed the formation of holes due to colonization by strain *Streptomyces* sp. SC-17 on the surface of PHB film.

**Figure 9.**
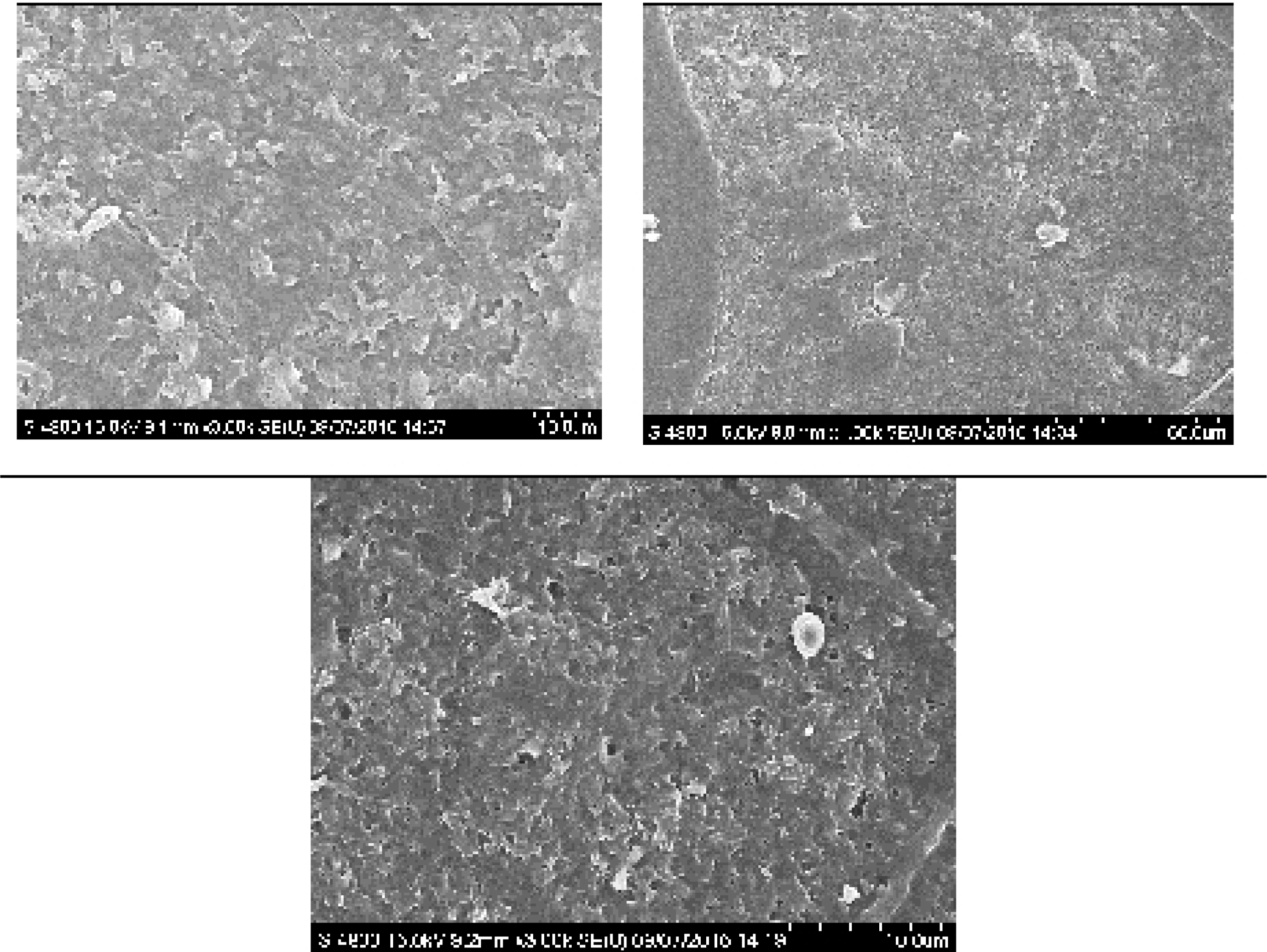
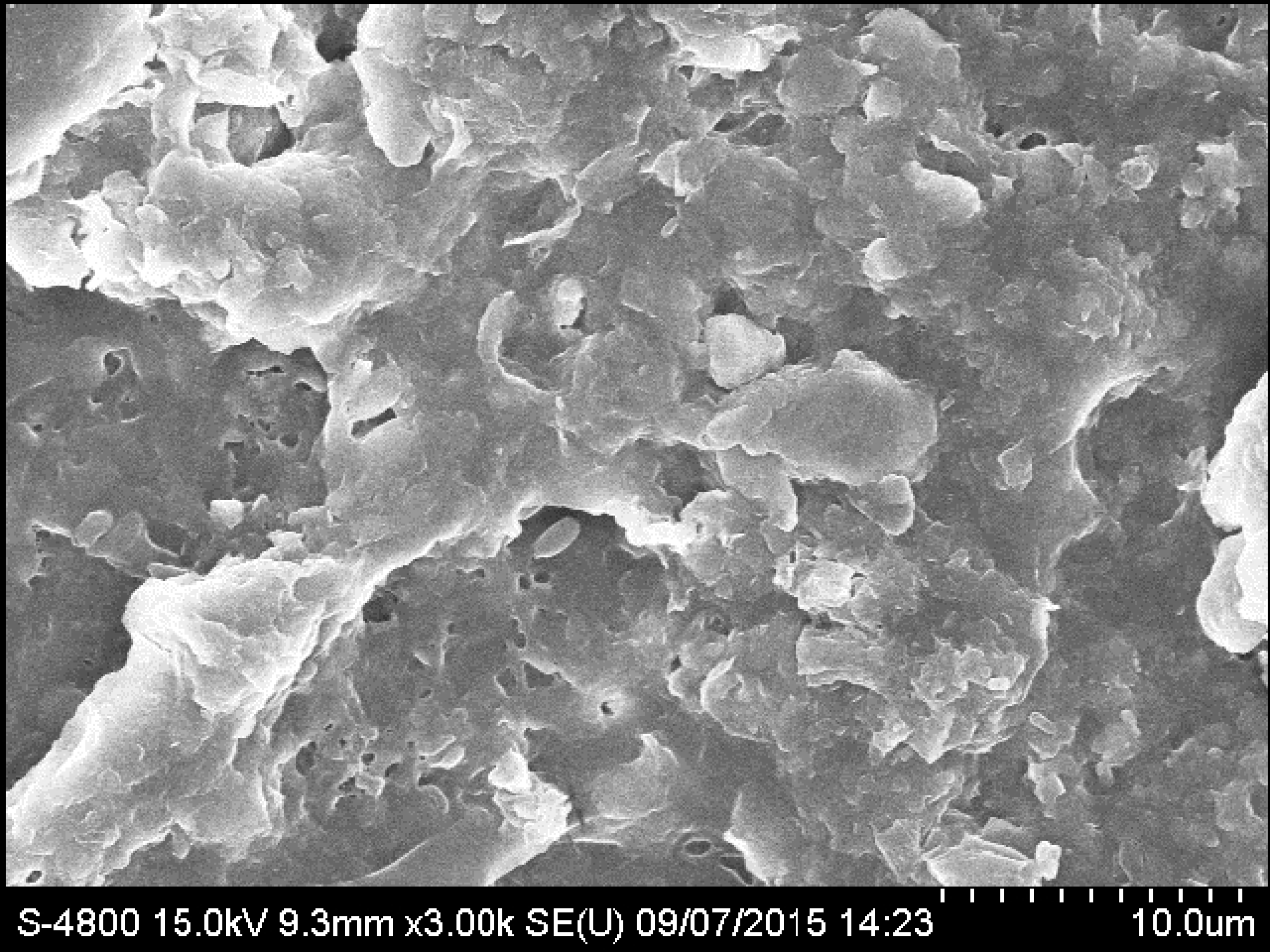
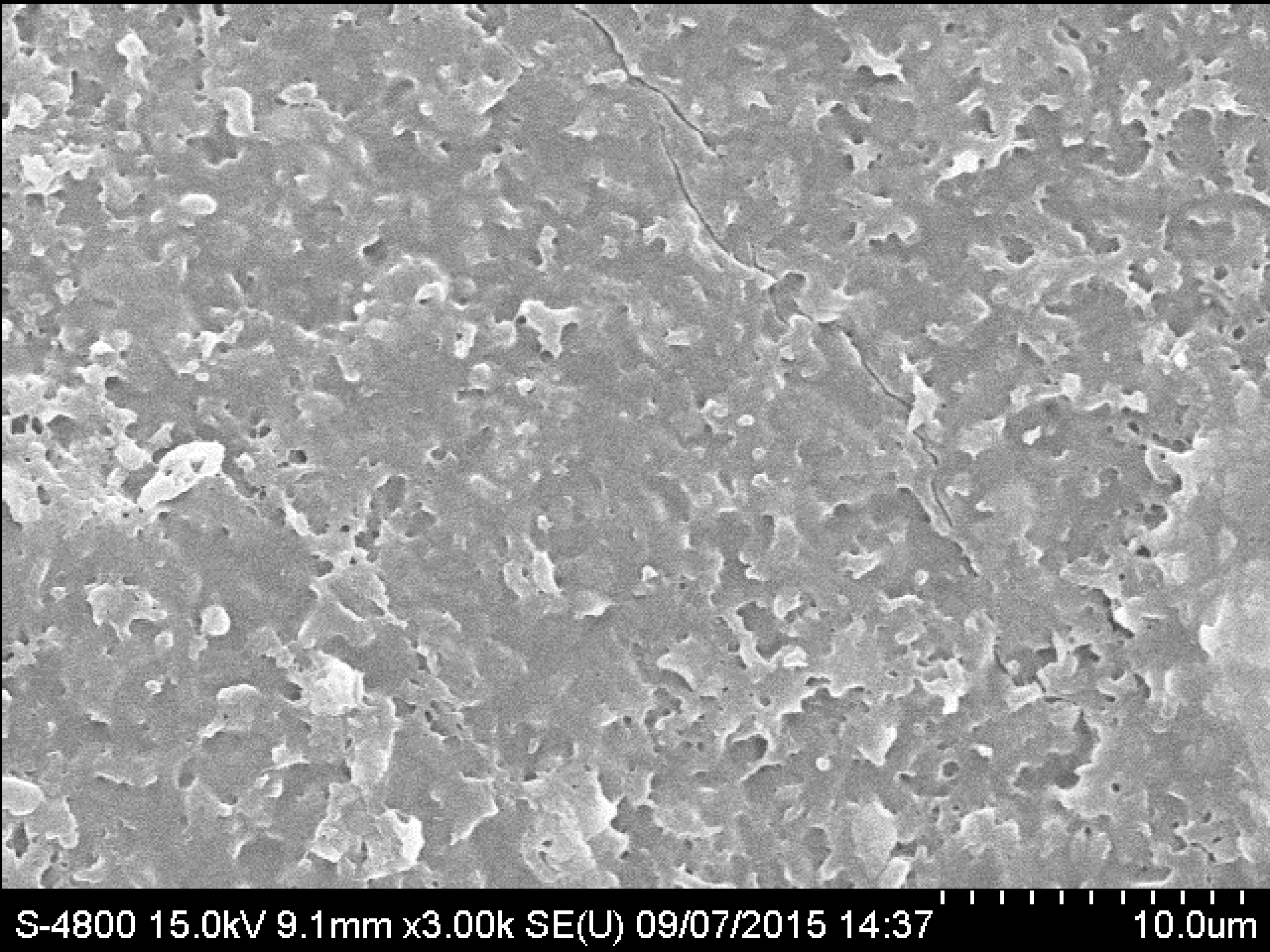
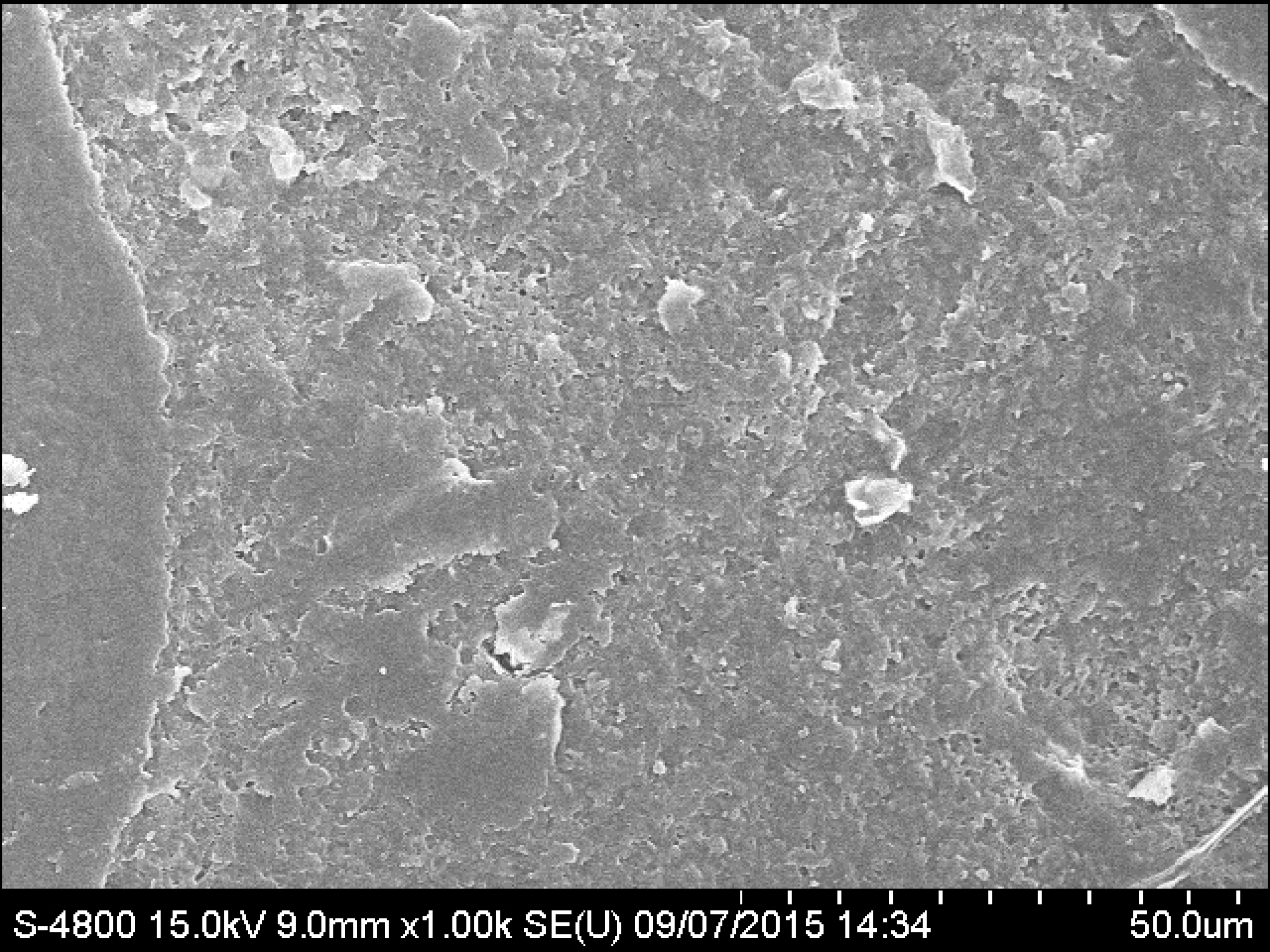
Changes in surface morphology of PHB film after soil burial experiment analyzed by FE SEM; (A) Control set, (B) PHB film buried in presence of natural soil, (C) PHB film degraded by *Stenotrophomonas* sp. RZS 7

## Conclusion

The present attempt provided PHB degrading *Stenotrophomonas* sp. RZS 7 which utilized PHB as a sole source of carbon under the influence of extracellular PHB depolymerase. The enzyme acted as a key enzyme responsible for biodegradation of PHB. Purification of PHB depolymerase by solvent purification resulted in protein precipitation and denaturation of the enzyme. Column chromatography by using octyl-Sepharose CL-4B came out as efficient and best purification method as it purification yielded maximum protein, maximum enzyme activity, and maximum specific activity. The molecular weight of purified PHB depolymerase of *Stenotrophomonas* sp. RZS 7 (40 kDa) matched with a molecular weight of *Aureobacterium saperdae*.

Biodegradation of PHB in liquid culture medium and under natural soil conditions confirmed PHB biodegradation potential of *Stenotrophomonas* sp. RZS 7. The results obtained in FTIR analysis, HPLC study and GC-MS analysis confirmed the biodegradation attempt in liquid medium by *Stenotrophomonas* sp. RZS 7. Changes in surface morphology of PHB film in soil burial as observed in FE SEM analysis confirmed the biodegradation of PHB. The isolate was capable of degrading PHB and resulted in 87.74% degradation. Higher rate of degradation under natural soil condition is the result of activity of soil microbes that complemented the degradation by *Stenotrophomonas* sp. RZS 7.

## Competing interests

All authors declare no conflict of interest.

## Acknowledgment

The authors would like to extend their sincere appreciation to the Deanship of Scientific Research at King Saud University, Saudi Arabia and MOE and UTM-RMC, HICOE, Malaysia for funding this research through the Research Group Project No. RG-1440-053 and R.J130000.7846.4J262.

## Data Availability Statement

All relevant data are within the paper.

